# Diagnostic value of circulating tumor DNA as an effective biomarker in cervical cancer: a meta-analysis

**DOI:** 10.1101/793869

**Authors:** Yulan Gu, Chuandan Wan, Jiaming Qiu, Yanhong Cui, Tingwang Jiang

**Author notes:** These authors contributed equally to this work (Yulan Gu, Chuandan Wan). Correspondence to: Chuandan Wan.

## Abstract

The applications of liquid biopsy have attracted much attention in biomedical research in recent years. Circulating cell-free DNA (cfDNA) in the serum may serve as a unique tumor marker in various types of cancer. Circulating tumor DNA (ctDNA) is a type of serum cfDNA found in patients with cancer and contains abundant information regarding tumor characteristics, highlighting its potential diagnostic value in the clinical setting. However, the diagnostic value of cfDNA as a biomarker in cervical cancer remains unclear. Here, we performed a meta-analysis to evaluate the applications of ctDNA as a biomarker in cervical cancer. A systematic literature search was performed using PubMed, Embase, and WANFANG MED ONLINE databases up to March 18, 2019. All literature was analyzed using Meta Disc 1.4 and STATA 14.0 software. Diagnostic measures of accuracy of ctDNA in cervical cancer were pooled and investigated. Fifteen studies comprising 1109 patients with cervical cancer met our inclusion criteria and were subjected to analysis. The pooled sensitivity and specificity were 0.52 (95% confidence interval [CI], 0.33–0.71) and 0.97 (95% CI, 0.91–0.99), respectively. The pooled positive likelihood ratio and negative likelihood ratio were 16.0 (95% CI, 5.5–46.4) and 0.50 (95% CI, 0.33–0.75), respectively. The diagnostic odds ratio was 32 (95% CI, 10–108), and the area under the summary receiver operating characteristic curve was 0.92 (95% CI, 0.90– 0.94). There was no significant publication bias observed. In the included studies, ctDNA showed clear diagnostic value for diagnosing and monitoring cervical cancer. Our meta-analysis suggested that detection of human papilloma virus ctDNA in patients with cervical cancer could be used as a noninvasive early dynamic biomarker of tumors, with high specificity and moderate sensitivity. Further large-scale prospective studies are required to validate the factors that may influence the accuracy of cervical cancer diagnosis and monitoring.

## Introduction

Human papillomavirus (HPV) is a type of papillomavirus that infects human skin and mucosa squamous epithelial cells. HPVs are DNA double-stranded spherical small viruses with a diameter of about 55 nm. The HPV genome contains approximately 7900 bases and can be divided into three functional regions [1]. The proteins E6 and E7, encoded by the early genes of HPV, can inhibit the functions of p53 and pRh in normal cervical epithelial cells and cause abnormal proliferation of cancerous cells, resulting in the development of genital warts and atypical proliferation of epithelial cells [2]. The immune system of most patients can eliminate HPV within approximately 9–16 months after infection. However, persistent infection by some high-risk HPVs, particularly HPV16 and HPV18, may lead to cervical cancer [3–5].

Cervical cancer is the fourth most common cancer among women worldwide. However, 85% of cases occur in developing countries [6]. Cervical cancer is now relatively uncommon in high-income countries owing to the introduction of HPV screening programs and HPV vaccines, which have led to a 70% decrease in the incidence and mortality rates of cervical cancer over past several decades [7]. Despite major advances in detection and prevention, an estimated 530,000 cases were recorded, and nearly 90% of 270,000 deaths occurred in middle- and low-income developing countries in 2012 [8]. There is still a need for minimally invasive and specific tests for HPV-induced cancer.

Recent progress in the analysis of blood samples for circulating tumor cells or cell-free circulating tumor DNA (ctDNA) has shown that liquid biopsies may have potential applications in the detection and monitoring of cancer [9–11]. Similarly, in a study on cervical cancer, ctDNA has become a major focus, providing a strong basis for early diagnosis and prognosis in cervical cancer [10, 12, 13]. Cervical cancer is typically caused by high-risk HPVs (hrHPVs), primarily genotypes 16 and 18 [4]. hrHPVs linearize DNA for integration into the cervical host genome and induce the expression of E6 and E7 genes, which are involved in the oncogenesis of cervical cancer [14, 15]. Cervical cancer cells and ctDNA harbor genomic rearrangements that can be released into the patient’s peripheral blood. From a diagnostic monitoring viewpoint, the consistent presence of HPV DNA in the blood of patients with cervical cancer can be used as a tumor marker. Although the mechanism mediating this phenomenon is unclear, the presence of such ctDNA in cervical cancer shows some diagnostic value. Interestingly, some studies have shown that circulating HPV DNA acts as a tumor DNA marker in patients with primary tumors caused by HPV infection [10]. Many recent studies have focused on ctDNA in cervical cancer; however, the exact relationships are still unclear [12, 16–18].

Accordingly, in this study, we performed a comprehensive analysis of the precise value of ctDNA for the diagnosis of cervical cancer.

## Materials and Methods

### Search strategy

This meta-analysis was conducted following the criteria of Preferred Reporting Items for Systematic Review and Meta Analyses [19]. A literature search was systematically performed using PubMed, Embase, Cochrane Library, and WANFANG medicine online databases for all relevant articles without language or regional limitations. No limitations were set with regard to the start date for publication, and the search ended on March 18, 2019. The following search terms were used: “cervical cancer AND ctDNA”, “cervix cancer AND ctDNA”, “cervical carcinoma AND ctDNA” OR “circulating DNA AND cervical cancer”. Various alterations in spelling and abbreviations were also used as search terms. Titles and abstracts were carefully screened for relevance, and duplicates were removed. The full text of each report that met the preliminary criteria was retrieved and assessed for inclusion into this meta-analysis.

### Inclusion and exclusion criteria

In this meta-analysis, eligible studies were selected according to these following inclusion criteria: (1) evaluated the diagnostic accuracy of quantitative analysis of ctDNA in cervical cancer; (2) the diagnostic value of ctDNA in cervical cancer was reported or could be calculated from the published data; (3) full text and all data could be retrieved and were available; (4) the techniques and target genes were clearly stated in the articles; (5) studies included at least 10 patients with cervical cancer and relevant negative controls. When the same patient population was used in several studies, only the most recent was included.

The exclusion criteria were as follows: (1) the diagnostic or prognostic value could not be deduced from incomplete data in the studies provided; (2) repeated studies from the same study group; (3) sample size less than 10; (4) data only from experiments based on cell lines; (5) studies published in languages other than English.

### Quality assessment

Two reviewers (CD Wan and YL Gu) independently reviewed and evaluated all eligible studies according to the Newcastle-Ottawa scale [20]. In case of disagreement, the decision was made by a third researcher, and disagreement was settled through discussion. The data extracted from the basic feature table included authors’ names, country, sample type, detection method, numbers of experimental and control groups, and analysis indicators. The outcome indicators included positives, false positives, false negatives, true negatives, sensitivity, and specificity. To assess the methodological quality of each study and potential risk of bias, QUADAS-2 Guidelines were used to evaluate the quality of all articles that met the inclusion criteria [21].

### Statistical analysis

We used standard methods recommended for meta-analysis of diagnostic test evaluations [19]. The meta-analysis was carried out with Meta-DiSc 1.4 and STATA 14.0 statistical software. The sensitivity was defined as the proportion of patients with ctDNA presence among all patients confirmed as having cervical cancer. The specificity was defined as the proportion of patients with negative ctDNA detection among all negative control volunteers without cervical cancer. The positive likelihood ratio (PLR) was calculated as sensitivity/(1 – specificity), whereas the negative likelihood ratio (NLR) was calculated as 1 – sensitivity/specificity. DOR was calculated as PLR / NLR and was used as an indication of how much greater the chance was of having cervical cancer for patients with ctDNA presence than for those without ctDNA. These indicators were summarized using a bivariate meta-analysis model, and the threshold effect was determined by receiver operative characteristic (ROC) curve and Spearman correlation analyses; *P* values of less than 0.05 indicated a significant threshold effect. Heterogeneity between studies was analyzed by chi-squared and *I*^*2*^ tests; a *P* value of less than 0.1 or an *I*^*2*^ higher than 50% indicated the existence of significant heterogeneity [22]. Meta-regression analysis was performed to explore the sources of heterogeneity. Deek’s funnel plot asymmetry test was used to test whether there was publication bias [23]. All statistical tests were two-sided, and results with *P* values of less than 0.05 were considered statistically significant.

## Results

### Study selection process

The initial search retrieved a total of 236 studies. As shown in Figure 1, 15 studies were eligible for review after carefully screening and rechecking. All relevant characteristics of these studies are summarized in Table 1. In total, 1109 patients with cervical cancer were evaluated in these studies published between 2001 and 2018. Among these studies, 11 enrolled patients from Asian countries/areas (one from Hong Kong, one from Thailand, one from Korea, one from India, one from Iran, two from Taiwan, and four from People’s Republic of China). Additionally, two studies were performed in France, and two were performed in America. Numerous review papers and duplicates between the literature databases were excluded.

**Table 1.** Main characteristics of all the studies enrolled the meta-analysis

| No. | Study | year | Patient number | Age(median and range) | region | method | Target gene | Sample source | Sample time | sensitivity | specificity |
| --- | --- | --- | --- | --- | --- | --- | --- | --- | --- | --- | --- |
| 1 | Pornthanakase m W[10] | 2001 | 50 | 50 | Thailand | qPCR | HPV DNA | plasma | BT | 36.00% | 100.00% |
| 2 | Dong SM[25] | 2002 | 232 | / | America | qPCR | HPV DNA | plasma | C | 48.70% | 98.33% |
| 3 | Hsu KF[35] | 2003 | 112 | 51(30-69) | Taiwan | qPCR | HPV DNA | serum | BT | 45.2% | 88.60% |
| 4 | Sathish N[36] | 2004 | 58 | 46.6(29-60) | India | PCR+RFLP | HPV DNA | plasma | BT | 48.2% | 100.00% |
| 5 | Yang HJ[27] | 2004 | 50 | 48(22-101) | HongKong | qPCR | HPV DNA | plasma | BT | 50% | 84.68% |
| 6 | Wei YC[37] | 2007 | 17 | 53.5(36-77) | Taiwan | Nested qPCR | HPV DNA | plasma | BT | 64.70% | 100.00% |
| 7 | Jaberipour M[38] | 2011 | 81 | 46.5(29-77) | Iran | qPCR | HPV DNA | plasma | BT | 23.5% | 90.91% |
| 8 | Campitelli M[24] | 2012 | 16 | / | France | DIPS-PCR | HPV DNA | serum | BT | 81.25% | 100.00% |
| 9 | Zhang X[29] | 2012 | 109 | 46(25-72) | China | RT-qPCR | Bmi-1 mRNA | plasma | BT | 69.70% | 95.90% |
| 10 | Jeannot E[32] | 2016 | 47 | / | France | ddPCR | HPV DNA | serum | BT | 83.00% | 100.00% |
| 11 | Zhang L[30] | 2016 | 89 | 46 | China | RT-qPCR | miR-21 | serum | BT | 78.00% | 80.00% |
| 12 | Zhang J[28] | 2017 | 168 | / | China | qMSP-PCR | MEG3 | plasma | UT | 90.47% | 79.80% |
| 13 | Kang Z[17] | 2017 | 21 | / | America | ddPCR | HPV DNA | serum | C | 90.48% | 100.00% |
| 14 | Kong Q[31] | 2017 | 18 | 39(23-61) | China | RT-qPCR | has-mir-92a | serum | BT | 69.60% | 80.40% |
| 15 | Kim HJ[26] | 2018 | 41 | / | Korea | qMSP-PCR | SIM1 | plasma | BT | 38.50% | 100% |

Sample time:BT, before treatment; C, combined; UT, undergoing treatment;

**Figure 1.**
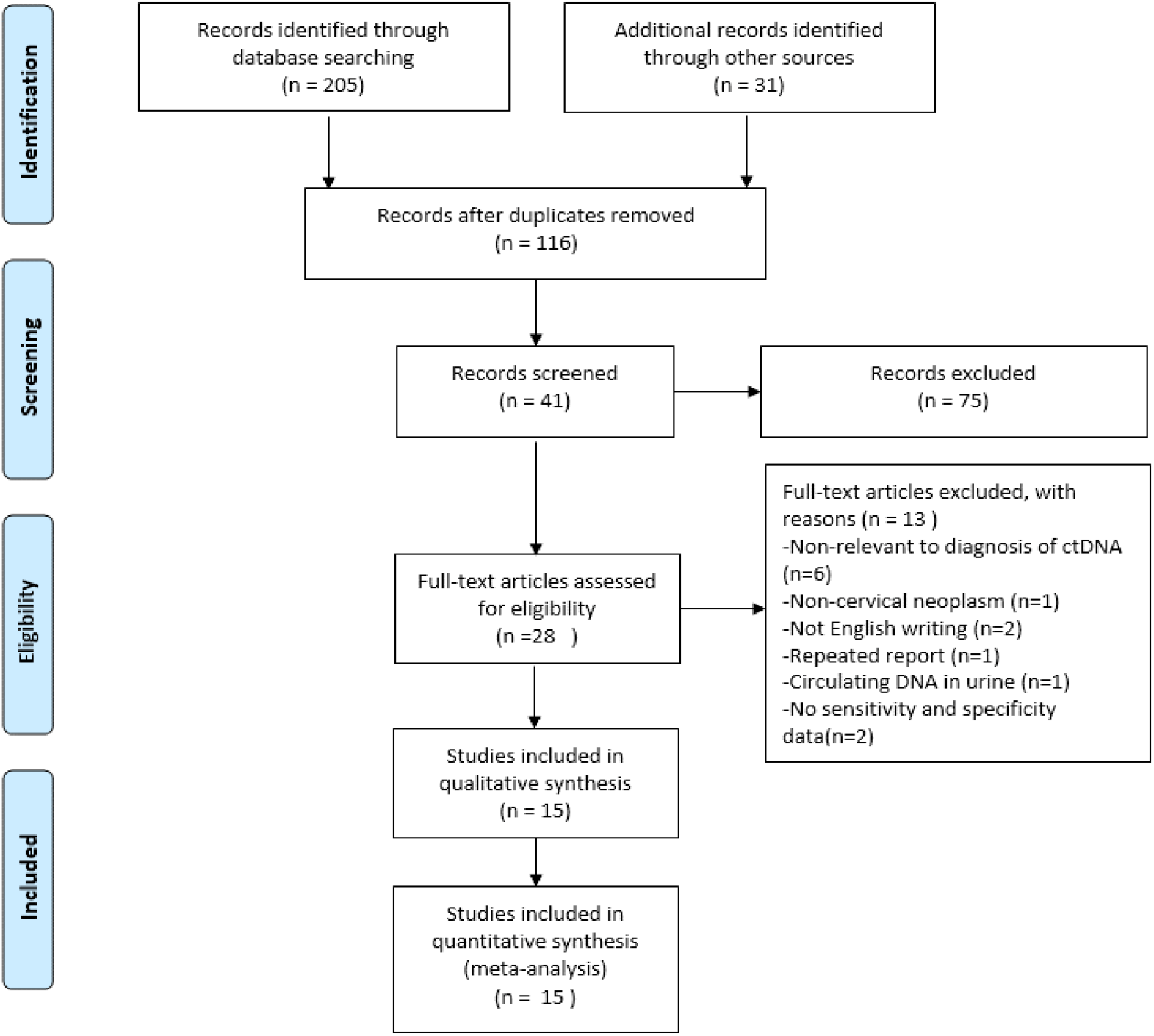
Flow chart of the enrolled studies.

### Review of eligible studies

The 15 eligible studies with data regarding the diagnostic value of ctDNA in cervical cancer are shown in Table 1. From these studies, 688 patients with cervical cancer were evaluated before treatment, and 421 patients were evaluated when undergoing treatment or after treatment. Patients with primary or metastatic cervical cancer with a TNM stage of I–IV who received surgery, chemotherapy, radiotherapy, or targeted therapy were included. The types of cervical cancer were squamous and adenomatous (approximate ratio of 4:1), as shown in Supplementary Table 1.

### Quality assessment

A quality assessment of the eligible studies was performed using QUADAS-2 (Figure 2). The included 15 studies were assessed using RevMan 5.3 software, and most of studies showed moderately low or unclear risk of bias. Two studies [10, 24] increased the risk of bias owing to a lack of patient selection. Four studies [25–28] did not mention the use of a blinding method or reference standard, which may have resulted in an unknown risk of bias in the meta-analysis.

**Figure 2.**
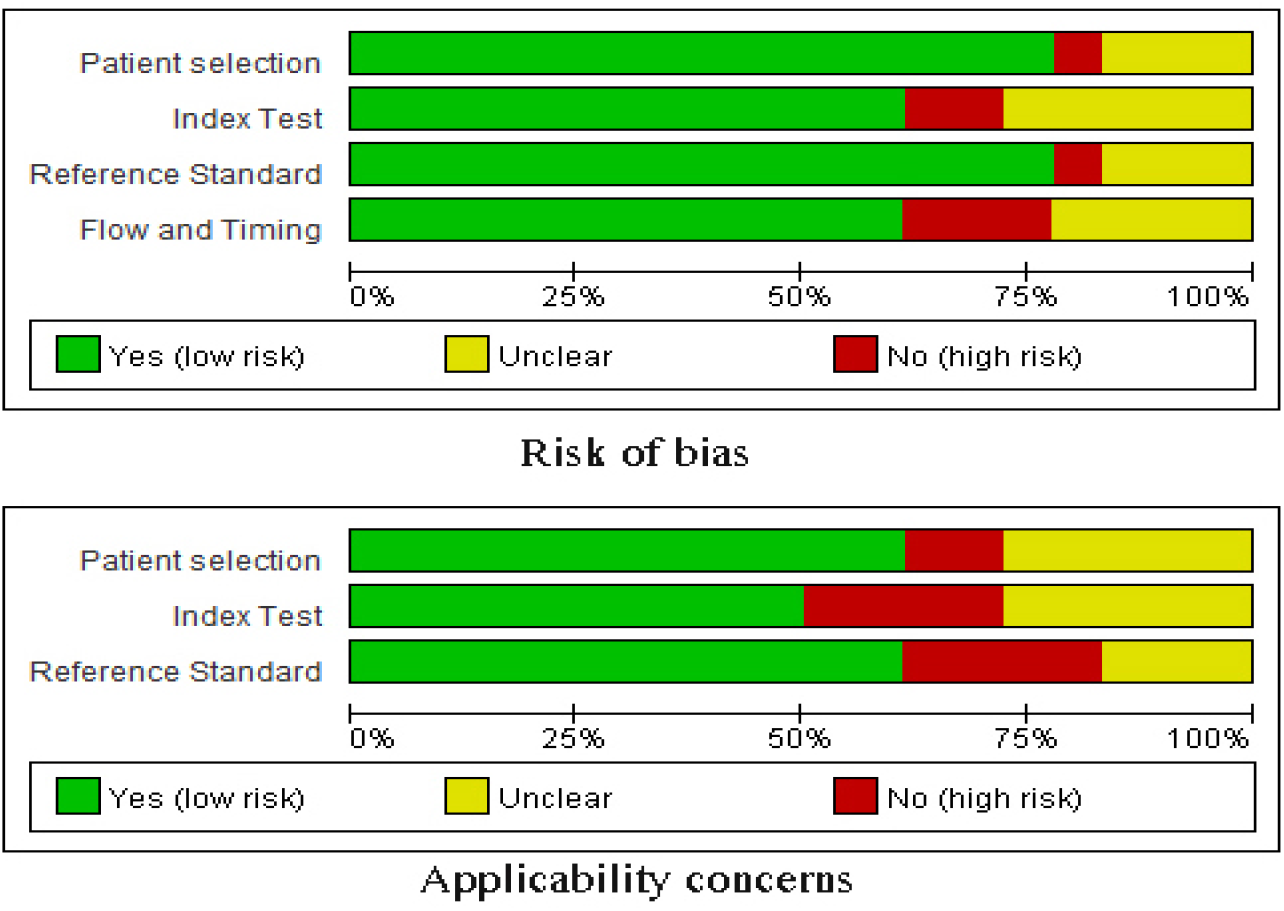
Quality assessment of the included studies according to QUADAS-2

### Detection of ctDNA and target genes

Polymerase chain reaction (PCR) was mainly applied to detect ctDNA in the studies included in this analysis. Two studies [10, 27] used Taqman PCR, and another three studies [29–31] used reverse transcription PCR. Additionally, two studies [17, 32] used droplet digital PCR (ddPCR), and three studies [24, 26, 28] used methylation-specific (MSP) PCR. Only one study [33] used nested PCR. Circulating HPV DNA or viral-cellular junction was detected in six studies [10, 17, 24, 27, 32, 33]. Other target genes were also different, such as *Bmi-1* mRNA [29], *miR-21* [30], *PIK3CA* mutations [34], *hsa-miR-92a* [31], and *SIM1* methylation [26]. Nine of the studies extracted ctDNA from plasma, and the other six studies extracted DNA from serum.

### ctDNA diagnostic accuracy in Cervical cancer

All of 15 studies were pooled into meta-analysis of diagnostic accuracy. The Spearman correlation coefficient was 0.594 (P> 0.05), suggesting that there was no threshold effect. Accordingly, heterogeneity owing to non-threshold effects was assessed with Q tests and I2 statistics. There was significant heterogeneity in the pooled sensitivity(I2= 97.2%, P<0.001) and specificity(I2= 87.1%, P< 0.001); Thus a random effects model would be applied to analyze the diagnostic parameters. As presented in Figure 3, the overall pooled sensitivity and specificity were 0.52(95% CI 0.33-0.71) and 0.97(95%CI 0.91-0.99), respectively. The overall pooled positive likelihood ratio (PLR) and negative likelihood ratio (NLR) were 16.0 (95%CI 5.5-46.4) and 0.50 (95%CI 0.33-0.75), respectively. The pooled diagnostic odd ratio (DOR) was 32 (95%CI 10-108). The summary receiver operator characteristic curve (SROC) was presented in Figure 4; the area under the SROC curve AUC was 0.92 (95%CI 0.90-0.94).

**Figure 3:**
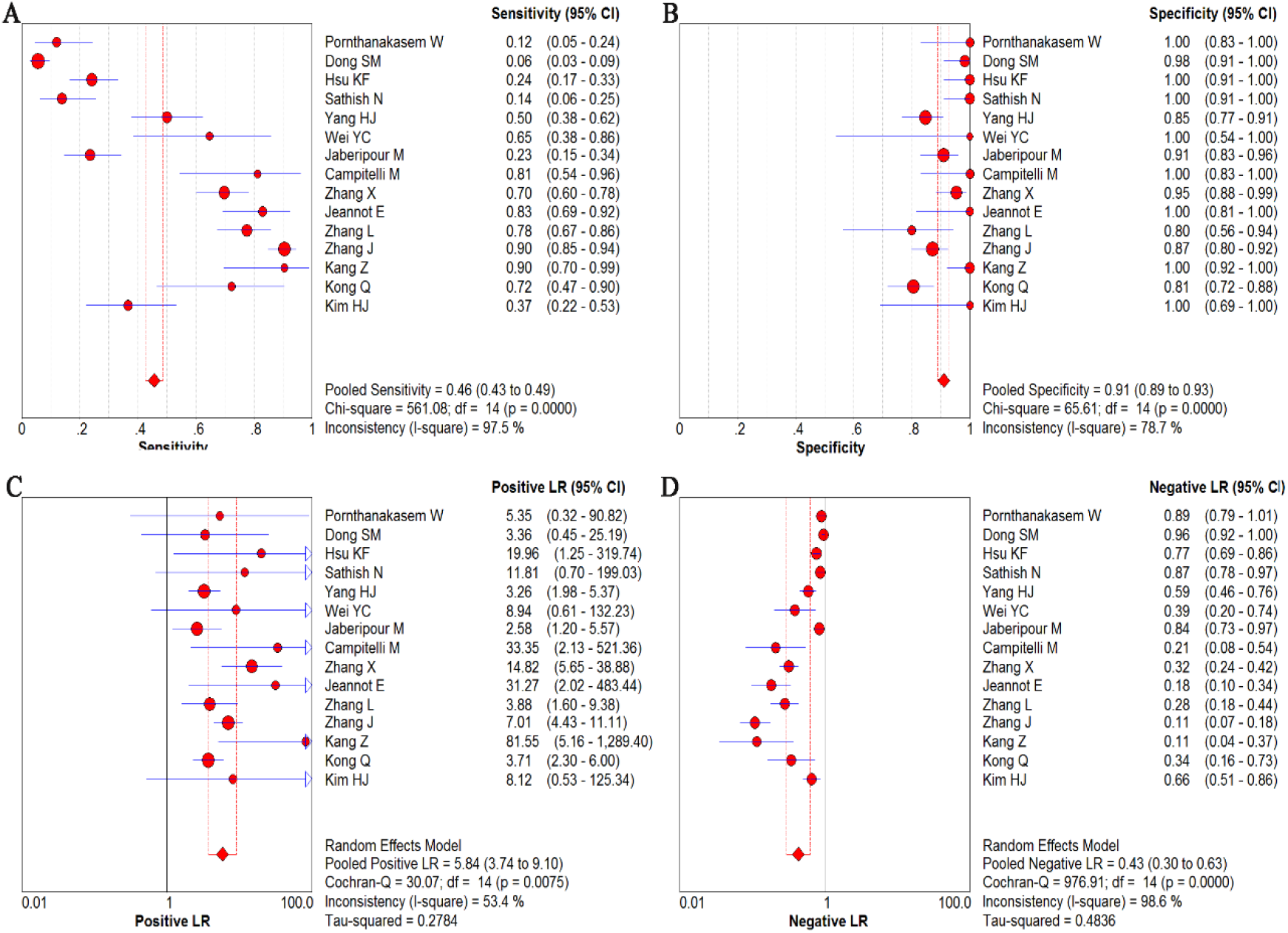
Diagnostic accuracy forest plots. (A) Forest plots of pooled sensitivity. (B) Forest plots of pooled specificity. (C) Forest plots of PLR. (D) Forest plots of NLR

**Figure 4:**
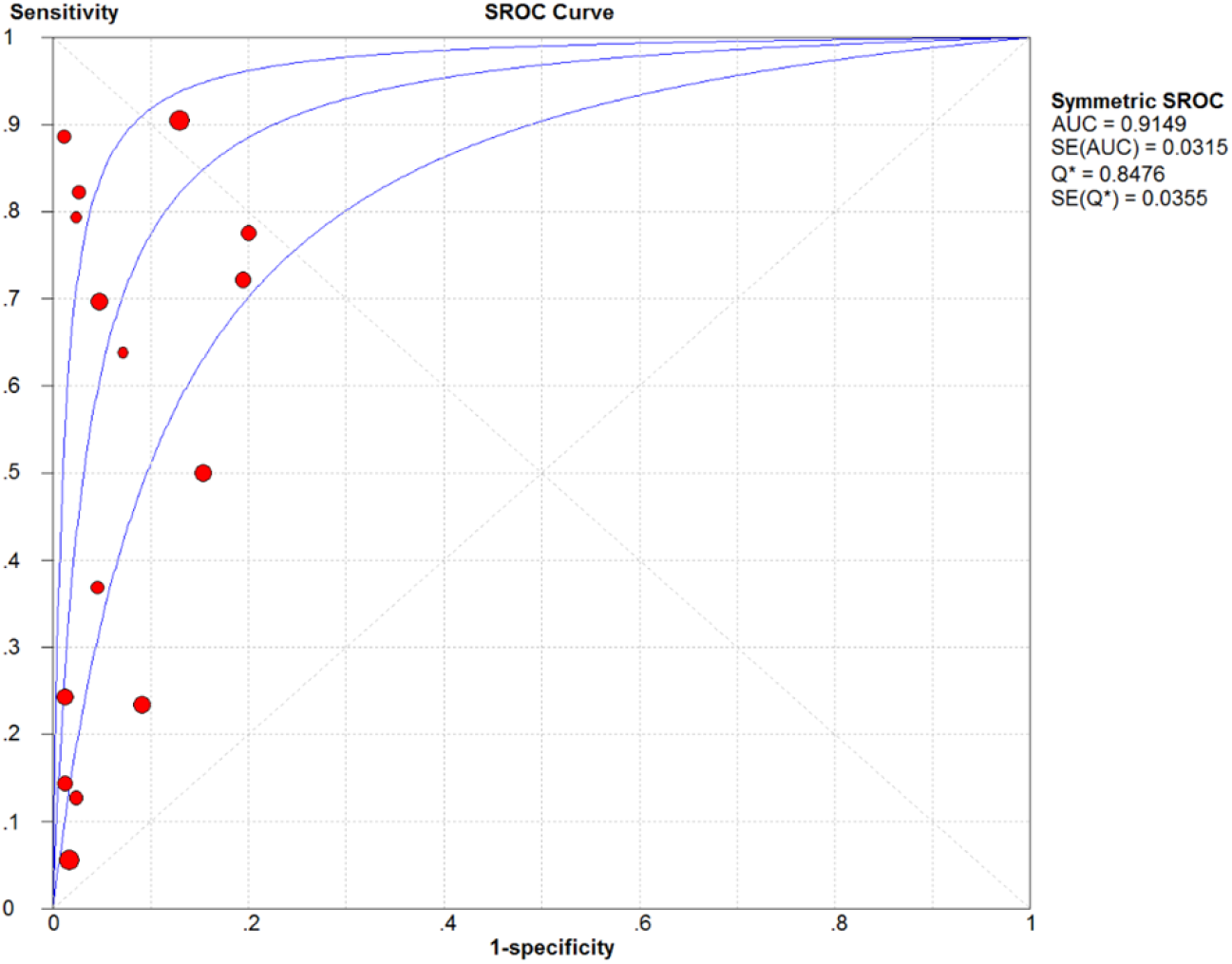
Summary receiver operating characteristic plot for the pooled studies diagnosis

### Subgroup analysis and meta-regression

Subgroup analysis was performed to explain the source of the significant heterogeneity in the diagnostic analysis. These different parameters in all of included studies were conducted including sample source (serum versus plasma), sample time (before treatment versus others), race (Asian versus Caucasian), gene target (HPV DNA versus others), and detection method (qPCR vs MSP and ddPCR). These diagnostic parameters of subgroups were showed in Table 2B. Meta-regression based on those five factors were applied to investigate the source of heterogeneity. However, none of these five factors showed any significantly influence on heterogeneity of universal diagnostic value (P>0.05) (Table 2A).

**Table 2A.** Meta regression of diagnostic value

| **parameter** | **Coef** | **SE** | **RDOR(95%CI)** | ***P*** |
| --- | --- | --- | --- | --- |
| Source | -0.746 | 1.115 | 0.470(0.030-6.620) | 0.525 |
| Method | -1.044 | 1.106 | 0.350(0.030-4.810) | 0.377 |
| Gene | -0.099 | 1.204 | 0.910(0.050-15.610) | 0.937 |
| Region | -1.082 | 1.735 | 0.340(0.010-20.500) | 0.553 |
| Time | -0.374 | 1.337 | 0.690(0.030-16.240) | 0.788 |
| Patient number | -0.044 | 0.943 | 0.960(0.100-8.890) | 0.964 |

**Table 2B.** Results of subgroups analysis

| Subgroup | Sensitivity(95% CI) | Specificity(95% CI) | PLR(95% CI) | NLR(95% CI) | DOR(95% CI) | AUC(95% CI) |
| --- | --- | --- | --- | --- | --- | --- |
| **Source** |  |  |  |  |  |  |
| Plasma | 0.46(0.43-0.49) | 0.91(0.89-0.93) | 5.84(3.74-9.10) | 0.43(0.30-0.63) | 18.70(8.53-41.00) | 0.91(0.85-0.98) |
| Serum | 0.59(0.54-0.65) | 0.90(0.86-0.93) | 10.91(3.22-37.01) | 0.27(0.09-0.83) | 38.97(11.08-137.12) | 0.86(0.78-0.93) |
| **Method** |  |  |  |  |  |  |
| qPCR | 0.35(0.31-0.38) | 0.91(0.89-0.93) | 5.05(3.04-8.40) | 0.56(0.41-0.75) | 12.00(5.58-25.80) | 0.87(0.78-0.97) |
| MSP and ddPCR | 0.81(0.75-0.85) | 0.91(0.85-0.95) | 7.62(4.90-11.85) | 0.23(0.06-0.96) | 64.67(32.92-12.705) | 0.95(0.92-0.97) |
| **Gene target** |  |  |  |  |  |  |
| HPV DNA | 0.27(0.24-0.30) | 0.94(0.92-0.96) | 6.85(3.09-15.21) | 0.60(0.46-0.78) | 15.25(5.42-42.94) | 0.95(0.85-0.99) |
| Others | 0.76(0.72-0.80) | 0.87(0.83-0.90) | 5.96(3.41-10.41) | 0.30(0.14-0.61) | 26.43(11.55-60.48) | 0.91(0.84-0.96) |
| **Region** |  |  |  |  |  |  |
| Mongolian | 0.53(0.50-0.57) | 0.89(0.87-0.92) | 4.96(3.33-7.37) | 0.49(0.35-0.70) | 14.02(6.18-31.79) | 0.89(0.82-0.96) |
| Caucasian | 0.27(0.22-0.32) | 0.99(0.96-1.00) | 18.65(4.04-86.13) | 0.26(0.01-7.97) | 76.54(5.92-989.35) | 0.98(0.97-0.99) |
| **Time** |  |  |  |  |  |  |
| Before treatment | 0.47(0.43-0.51) | 0.90(0.88-0.93) | 5.37(3.25-8.89) | 0.51(0.39-0.67) | 13.83(6.67-28.66) | 0.87(0.77-0.97) |
| Under- or after treatment | 0.44(0.39-0.49) | 0.93(0.88-0.96) | 8.61(2.46-30.05) | 0.23(0.00-121.78) | 45.22(4.09-499.96) | 0.95(0.92-0.97) |
| **Patient Number** |  |  |  |  |  |  |
| <50 case | 0.54(0.48-0.60) | 0.89(0.85-0.92) | 6.92(3.05-15.69) | 0.38(0.21-0.67) | 23.44(7.27-75.58) | 0.91(0.76-0.99) |
| ≥50 case | 0.43(0.40-0.46) | 0.93(0.90-0.95) | 5.82(3.35-10.11) | 0.49(0.29-0.83) | 15.42(4.84-49.08) | 0.92(0.85-0.99) |
| Overall | 0.46(0.43-0.49) | 0.91(0.89-0.93) | 5.84(3.74-9.10) | 0.43(0.30-0.63) | 18.70(8.53-41.00) | 0.91(0.85-0.98) |

### Sensitivity analysis

To further explore the heterogeneity of the included studies, a sensitivity analysis was conducted by removing individual studies. As shown in Figure 3, no outlier study was identified, and the results were considerable stable and reliable.

**Figure 5.**
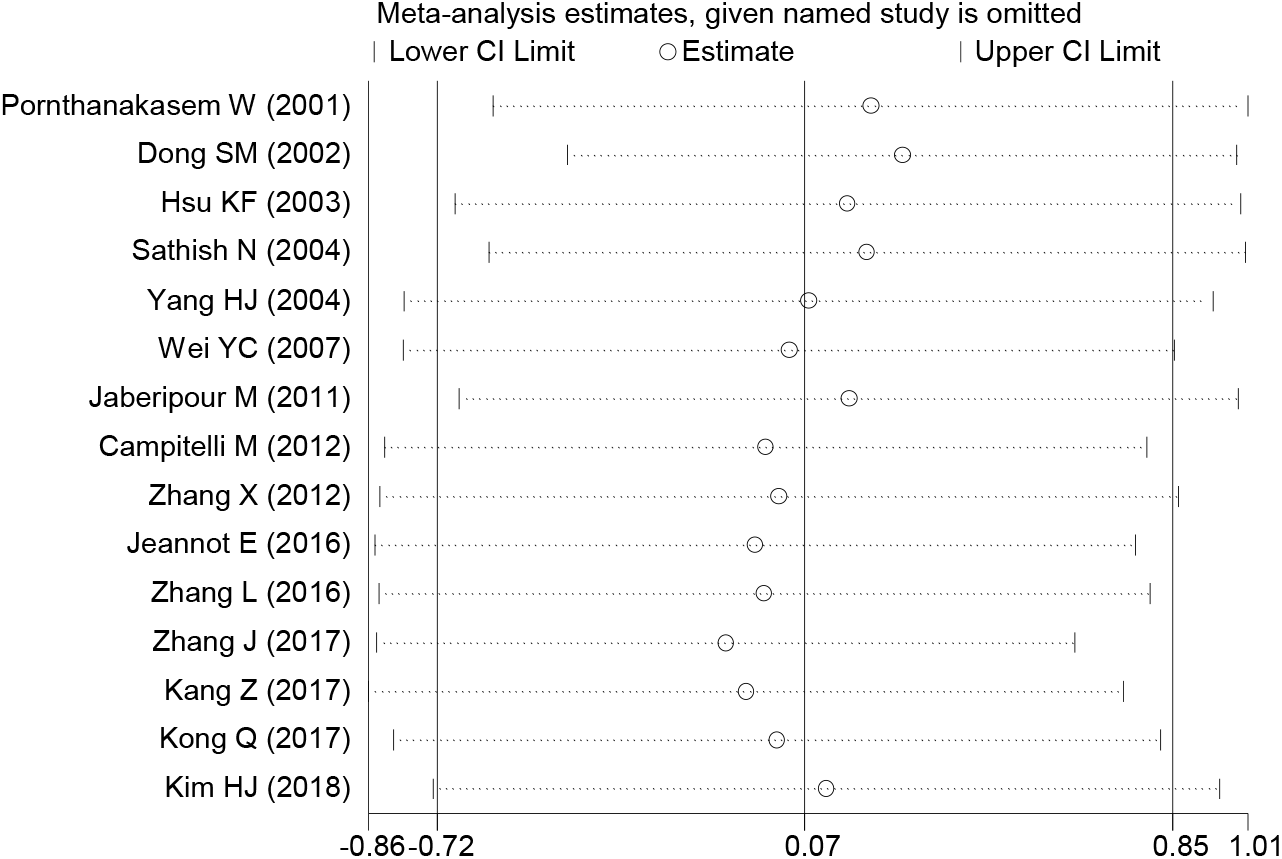
Sensitivity analysis of the overall pooled study.

### Publication bias

We applied Deeks’ funnel plot asymmetry tests to estimate the publication bias of the included studies. As shown in Figure 4, the regression line was nearly vertical, confirming the lack of significant publication bias across the overall enrolled studies (*P* = 0.54).

**Figure 6.**
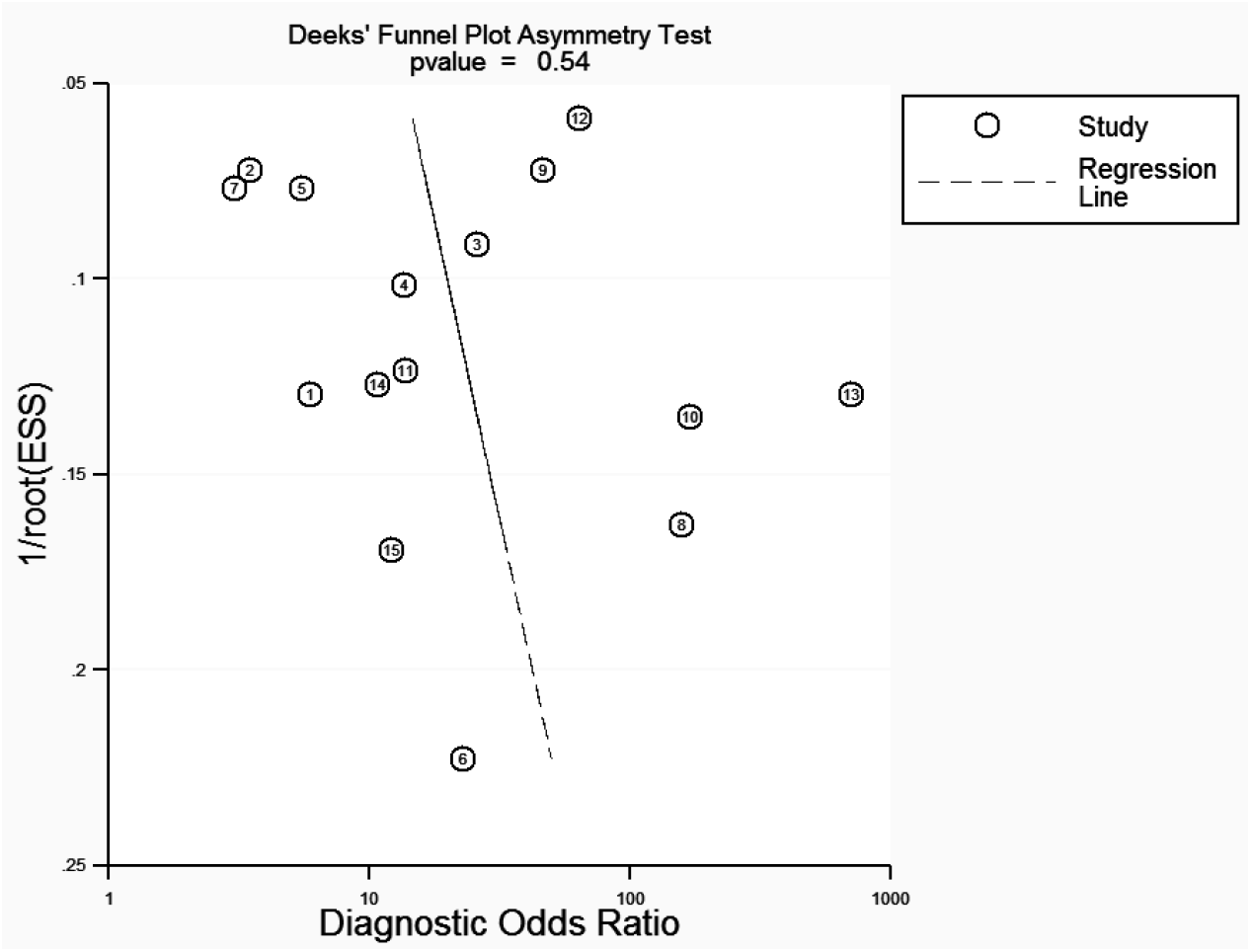
Deek’s funnel plot to assess publication bias. ESS, effective sample size.

## Discussion

In pathological analysis, traditional surgical specimens are used as the gold standard for obtaining fundamental information for clinical decision-making. However, obtaining such specimen directly from tumors is an invasive procedure and can delay the observation of tumor dynamic changes after treatment [39]. Cancers are known to shed tumor cell DNA into the blood stream [34], and examining the levels and mutations in ctDNA can provide almost real-time information regarding tumor status. Once a patient achieves remission, liquid biopsy has the potential to improve post-treatment surveillance by following subtle changes in tumor cfDNA. These changes may indicate disease recurrence prior to clinical manifestation. Therefore, liquid biopsy has recently been extensively investigated as a potential new diagnostic technique [34, 40].

Despite major advances in early detection, including Pap smears and co-human papillomavirus testing, there are still an estimated 5 million new cases of cervical cancer and 2 million cervical cancer related deaths worldwide, making this disease the fourth leading cause of cancer-related death in women [41]. Accordingly, there is a need for a minimally invasive and specific test for disease monitoring, which could be beneficial for patients with cervical cancer. Circulating cfDNA has been widely evaluated using liquid biopsies for detecting cancer, monitoring disease, characterizing drug targets, and uncovering resistance in various tumors [40, 42–44].

Many previous meta-analyses have reported that the diagnostic accuracy of quantitative analysis of ctDNA is superior to conventional biomarkers for the diagnosis of several cancers, including ovarian cancer [22], lung cancer [45], gastric cancer [39], and colon cancer [46]. However, to the best of our knowledge, this is the first meta-analysis exploring ctDNA in patients with cervical cancer. Meta-analyses are powerful tools for summarizing the results from different and related studies by producing a single estimate of the major effects with enhanced precision. This approach can overcome the problem of small sample size and inadequate statistical power in genetic studies of complex traits and provide more reliable results than single case-control studies [47]. Because the relationship between ctDNA and cervical cancer is still unclear, we performed a comprehensive analysis of the clinical utility of ctDNA in the diagnosis of patients with cervical cancer.

This meta-analysis combined the outcomes of 1109 patients with cervical cancer from 15 individual studies, investigating the diagnostic values of six nucleotide targets, including HPV DNA, *MEG3*, *SIM1*, *Bmi* mRNA, and *miR-21*. From the 15 studies, the pooled sensitivity and specificity were 0.52 (95% CI, 0.33–0.71) and 0.97 (95% CI, 0.91–0.99), respectively. The LR reflects the characteristics of sensitivity and specificity and is not affected by the prevalence rate. Therefore, LR is more stable than sensitivity and specificity. LRs of greater than 10 or less than 0.1 indicate large and often conclusive shifts from pretest to post-test probability [48]. In this meta-analysis, the overall pooled PLR and NLR were 16.0 (95% CI, 5.5–46.4) and 0.50 (95% CI, 0.33–0.75), respectively. This result indicated that patients with cervical cancer had approximately 16 times greater chance of being ctDNA positive than normal controls, with an error rate of approximately 50% when the true negative was determined in the ctDNA negative test. The pooled DOR was 32 (95% CI, 10–108), which indicated a relatively high accuracy of ctDNA in cervical cancer. Summary ROC (SROC) can be applied to summarize overall test performance, and the area under the SROC curve (AUC) was 0.92 (95% CI, 0.90–0.94), suggesting that ctDNA in the plasma or serum of patients with cervical cancer had excellent accuracy for diagnosing cervical cancer. Because significant heterogeneity existed, if relatively accurate diagnostic parameters were achieved, subgroup analysis would be needed to analyze the source. Subgroup analyses revealed that the heterogeneity of sensitivity could be related to the source of the specimen (e.g., plasma versus serum), the gene target (e.g., HPV DNA versus others), the region (e.g., Mongolian versus Caucasian), the time of specimen collection (e.g., before treatment versus after treatment), the number of patients (e.g., less than 50 versus greater than or equal to 50), and the method of analysis (e.g., quantitative PCR versus MSP-PCR and ddPCR). Ten of 15 included studies had selected circulating HPV DNA as a detected gene target. Other gene targets had higher sensitivity but lower specificity, including *SIM1* methylation [26], *MEG3* methylation [28], *Bmi-1* [29], *miR-21* [30], and *hsa-miR-92a* [31]. Most studies employed a qPCR method that demonstrated relatively high sensitivity and specificity. Over time, more accurate diagnostic parameters were obtained by ddPCR. We found that MSP-PCR and ddPCR were more accurate for detecting ctDNA in patients with cervical cancer than qPCR. In particular, the application of ddPCR in liquid biopsy greatly improves the diagnostic value of ctDNA in cervical cancer [40]. However, statistical regression data showed that all these differences between subgroups were not statistically significant (*P* > 0.05). Taken together, these results indicated that the study design did not substantially affect the diagnostic accuracy. Heterogeneity may have been caused by other factors, such as patient age, tumor type, tumor size, TNM stage, and differences in the experimental protocols, which could not be analyzed in the current study because of loss of data or unrecognizable details. Therefore, further studies with large sample sizes and more details, e.g., race, specimen features, and tumor properties, are needed to confirm these findings.

Cervical cancer differs from other cancers because HPV infection is a crucial step in tumorigenesis, accounting for 99.7% of cervical cancer cases. HPV16 and HPV18 are the two most important serotypes, identified in more than 70% of cervical carcinomas worldwide [49]. Circulating nucleic acids were first reported in the 1940s and included DNA and RNA with genomic, mitochondrial, and viral origins. Apoptotic cells, necrosis of tumor cells, and some living cells may actively release DNA fragments into the circulation. These specific changes in tumor DNA with regard to oncogenes, tumor-suppressor mutations, microsatellite alterations, and hypermethylation can be similarly detected in ctDNA. Detection of these specific changes in ctDNA may yield tumor markers with very high specificity, although high test sensitivity is also needed because mutant DNA fragments may represent a small proportion of total ctDNA. Although the 15 studies included in this meta-analysis had very high specificity, there was uneven sensitivity. The pooled results indicated that there was significant heterogeneity in sensitivity that could impact diagnostic accuracy. The Spearman correlation coefficient was 0.594 (*P* > 0.05), suggesting that the threshold effect was not the source of heterogeneity. Because the size of ctDNA fragments is generally approximately 200 bp [50], PCR primer pairs that target shorter DNA fragments may identify more patients with detectable HPV DNA. With more primer pairs, further increases in detection rates may be possible. Other influencing factors, such as patient number, specimen extraction time, region, and specimen source, may also influence these parameters; however, these differences were not statistically significant. In addition, publication bias was also not significant, indicating that the results of this meta-analysis were reliable and credible.

There were several limitations to this meta-analysis. First, the sensitivities of the included studies varied widely. Different gene detection methods and gene targets could have led to major differences. Therefore, significant heterogeneity between studies could not be avoided. The unique characteristics of ctDNA limit its sensitivity as a diagnostic indicator, and more sensitive and accurate detection techniques may need to be applied. Although subgroup and regression analyses were performed to explore the sources of heterogeneity, the results of these analyses explained few effectors. Second, some studies with limited patient numbers and controls were included in this meta-analysis, reducing the effectiveness of the combined statistical analysis. Relatively few papers on ctDNA in patients with cervical cancer have been published, and studies that met the inclusion criteria should not be excluded. Otherwise, serious publication bias cannot be avoided. Third, owing to the nature of our research, selected bias and incomplete searches could have occurred. Fourth, most of the included studies came from Asian countries. Although no studies have shown that genetic factors differ among different ethnicities in patients with cervical cancer, the physiological responses of patients with different ethnicities may lead to variations in ctDNA metabolism. Finally, different primers may have been used for PCR of same gene targets, resulting in a potential source of bias.

## Conclusions

Despite some limitations, this meta-analysis clearly indicated that ctDNA detection may be a very specific, but relatively sensitive test in patients with cervical cancer. Our findings provided reliable evidence that ctDNA was a promising potential biomarker for the diagnosis of cervical cancer. Of course, to obtain a more accurate statistical data analysis, additional studies with larger sample sizes from patients of different ethnicities will be necessary in the future.

## Acknowledgments

There was no funding source associated with this meta-analysis.

